# Neo-sex chromosome evolution shapes sex-dependent asymmetrical introgression barrier

**DOI:** 10.1101/2021.07.13.452191

**Authors:** Silu Wang, Matthew J. Nalley, Kamalakar Chatla, Reema Aldaimalani, Ailene MacPherson, Kevin Wei, Russ Corbett, Dat Mai, Doris Bachtrog

**Affiliations:** Integrative Biology, University of California, Berkeley, 3040 VLSB, Berkeley, CA, USA 94720; University of Toronto; Biomolecular Engineering and Genomics Institute, University of California, Santa Cruz

**Keywords:** Introgression, neo-sex chromosome, hybridization, Y degeneration, inversion, meiotic drive

## Abstract

It is increasingly recognized that sex chromosomes are not only the “battlegrounds” between sexes, but also the “Great Walls” fencing-off introgression between diverging lineages. Here we describe conflicting roles of nascent sex chromosomes on patterns of introgression in an experimental hybrid swarm. *Drosophila nasuta* and *D. albomicans* are recently diverged, fully fertile sister species that have different sex chromosome systems. The fusion between an autosome (Muller CD) with the ancestral X and Y gave rise to neo-sex chromosomes in *D. albomicans*, while Muller CDs remains unfused in *D. nasuta*. We found that a large block containing overlapping inversions on the neo-sex chromosome stood out as the strongest barrier to introgression. Intriguingly, the neo-sex chromosome introgression barrier is asymmetrical in a sex-dependent manner. Female hybrids showed significant *D. albomicans* biased introgression on Muller CD (neo-X excess), while males showed heterosis with excessive (neo-X, *D. nasuta* Muller CD) genotypes. While the neo-Y is a more compatible pairing partner of the neo-X, it also shows moderate levels of degeneration and may thus be selectively disfavored, and sex ratio assay revealed heterospecific meiotic drive. We used a population genetic model to dissect the interplay of sex chromosome drive, heterospecific pairing incompatibility between the neo-sex chromosomes and unfused Muller CD, neo-Y disadvantage, and neo-X advantage in generating the observed neo-X excess in females and heterozygous (neo-X, *D. nasuta* Muller CD) genotypes in males. We show that moderate neo-Y disadvantage and *D. albomicans* specific meiotic drive are required to counteract the effect of heterospecific meiotic drive observed in our cross, in concert with pairing incompatibility and neo-X advantage to explain observed genotype frequencies. Together, this hybrid swarm between a young species pair shed light onto the dual roles of neo-sex chromosome evolution in creating a sex-dependent asymmetrical introgression barrier at species boundary.

## Introduction

Speciation is a fundamental process that generates much of the diversity of life (1), yet the underlying genomic mechanisms of speciation are not well-understood (2–5). There has been accruing evidence of sex chromosomes disproportionately accumulating genomic differentiation among diverging lineages (6–9). However, the interaction of sex chromosome evolution and genomic differentiation underlying early speciation remains an open question (10). That is, what is the effect of chromosome evolution and differentiation on the extent and direction of introgression across species boundaries? The ideal system to investigate this question would encompasses lineages in the early stage of speciation, in which newly formed sex chromosomes evolve as speciation unfolds (11, 12).

The sister species *Drosophila albomicans* (distributed from Japan, China to Northeast India) and *D. nasuta* (found in East Africa, Sri Lanka and the India subcontinent) diverged around 0.15-0.5 mya (10, 11). These species are indistinguishable morphologically and show little to no pre-mating isolation (16) and only weak hybrid breakdown in advanced generation hybrids (17, 18), but have distinct sex chromosome configurations. *D. nasuta* harbors the ancestral genotype of this species group (2n = 8), while *D. albomicans* has a neo-sex chromosome pair, formed by the fusion of an autosome (Muller CD) and the ancestral sex chromosomes (Muller A) around 0.12 mya (19)(15, 20) (**Fig 1A**). Genetic studies have suggested that the neo-sex chromosomes evolved sequentially, with the X/ Muller-CD fusion (the neo-X) being selectively favored over the ancestral unfused chromosomes, and subsequently driving the fixation of the Y/Muller-CD fusion (the neo-Y chromosome) to overcome meiotic structural incompatibilities (21). Thus, this species pair provides a unique opportunity to investigate the role of sex chromosome evolution in speciation.

**Fig. 1.**
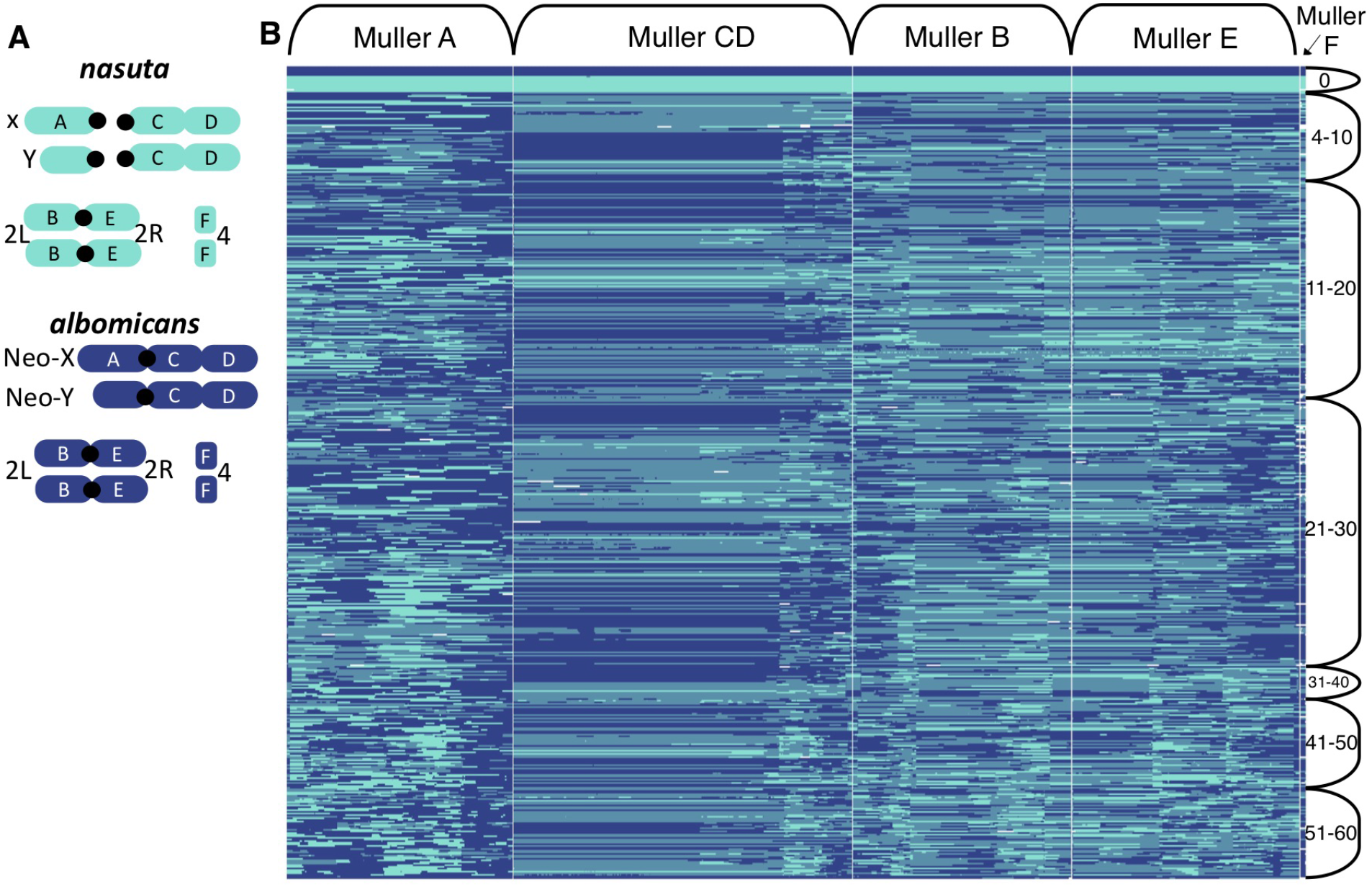
Genotypes and admixture between *D. albomicans* and *D. nasuta* ancestry. **A**, Karyotype of *D. albomicans* and *D. nasuta*. Muller CD and Muller A are separated in *D. nasuta* but fused in *D. albomicans*, forming a neo-X or neo-Y chromosome. **B**, Ancestry-HMM of haplotypes (in columns) in hybrids of various generations (rows, numbered on the right). The turquoise and royal blue represents homozygous *D. nasuta* and *D. albomicans* genotype, respectively and the heterozygous genotypes are represented by pale blue.

Here we investigated the role of the neo-sex chromosomes on patterns of genomic differentiation over more than 62 generations of hybridizations between these recently diverged sister species. Hybrid swarms undergoing hybridization over multiple generations can reveal the extent and direction of introgression in different parts of the genome, and allow us to investigate the role of sex chromosomes contributing to genomic differentiation. Specifically, tracking ancestry turnover in the neo-sex chromosome relative to the rest of the genome can reveal the role of sex chromosome evolution in shaping patterns of introgression.

The evolution of neo-sex chromosomes could shape introgression between *D. nasuta* and *D. albomicans* multiple ways. The fused neo-X could be selectively favored over the unfused *D. nasuta* karyotype, as suggested by hybrid crosses (21)(22); this could facilitate the spread of the neo-X in the hybrid population. Increased pairing compatibility of the neo-X and neo-Y during male meiosis could indirectly facilitate neo-Y introgression at high frequency of the neo-X (21). On the other hand, the neo-X could also limit or promote the spread of the neo-Y due to meiotic drive. Notably, a polymorphic sex ratio drive was discovered in crosses between *D. albomicans* strains from Japan, and *D. nasuta* strains from India (23), and in interspecific crosses between *D. albomicans* and *D. nasuta* (24). Thus, the neo-X may harbor meiotic drive alleles that are toxic to neo-Y carriers in certain *D. albomicans* strains (23, 24), and (some) neo-Y chromosomes may harbor suppressors for drive that are absent in *D. nasuta*, which would limit or promote the spread of the neo-Y, respectively. In addition, accumulation of frame-shift mutations and lower gene expression was observed for a subset of neo-Y genes (19). Beginning degeneration of the neo-Y incurs a fitness disadvantage over the unfused *D. nasuta* homologous Muller-CD chromosomes, and could select against the neo-Y chromosome in male hybrids. Altogether, the *D. nasuta* / *D. albomicans* system provides an opportunity to uncover and dissect the conflicting roles of neo-sex chromosome evolution in introgression.

Although this species pair are predominantly allopatric and typically do not form secondary contact in nature, artificial hybrid populations provide a valuable opportunity to investigate the effect of neo-sex chromosome evolution on patterns of introgression between these otherwise similar lineages. Here we generated replicate hybrid swarms of *D. nasuta* and *D. albomicans* and sequenced almost 450 sampled hybrid individuals over 62 generations. In particular, we ask: (1) what is the extent and direction of introgression at the species boundary; (2) does the direction and extent of introgression within neo-sex chromosome differ from the rest of the genome; (3) do sexes differ in introgression? We address these questions accounting for chromosomal inversions, neo-X advantage, neo-Y degeneration disadvantage, neo-X meiotic drive, and heterospecific pairing incompatibility.

## Methods

### Hybrid swarm and sampling

We used *D. nasuta* strain 15112-1781.00 (from Mysore, India) and *D. albomicans* strain 15112-1751.03 (from Nankang, Taiwan) to construct admixed populations (‘hybrid swarms’). These strains have chromosome-level genome assemblies and differ by two fixed inversions on Muller-CD (https://doi.org/10.1101/2021.06.01.446624). We set up reciprocal interspecific crosses (i.e. we crossed 30 *D. nasuta* virgin females with 30 *D. albomicans* males; and 30 *D. abomicans* virgin females with 30 *D. nasuta* males), and mixed all of the resulting F1 offspring to initiate the hybrid swarm. The hybrid swarm was maintained in large Plexiglass population cages (dimension being 12” x 12” x 12”). The population cages were kept at humidity 48%, and a 12h light-dark cycle with lights on during 8AM-8PM. Each week, we added two new molasses bottles of fly food (standard corn medium); these bottles were removed from the cages 4 weeks later, all adults in the bottles were discarded, and newly emerging flies from the sample bottles were collected and frozen about one week later. We set up two independent replicates of the hybrid swarm (one in 2014, and one in 2018). We also set up an additional population cage, where we simply combined roughly 100 adults from both species to initiate the hybrid swarm; this cage was maintained and sampled just like the F1 hybrid cages. Generations of sampled flies were determined based on sampling date, assuming a generation time of 14 days.

### DNA isolation, library preparation and sequencing

We sequenced a total of 232 females and 215 males sampled from select generations (between 3 to 62; **Table S1**). The detailed sample size in terms of sex and generation is summarized in **Table S1**. DNA extractions were performed as described in (25) with modifications as follows. Flies were crushed in Puregene lysis buffer (120 ul) using the Mixer Mill 400 at 30 Hz for 3 minutes. Lysate (100 ul) was treated with RNase A, and protein was precipitated with Puregene protein precipitation solution (33 ul) on ice for 30 minutes. Clarified lysate (80 ul) was transferred to ice cold isopropanol (80 ul), mixed well, and incubated for 30 minutes. DNA was precipitated at maximum speed for 30 minutes. Pellets were washed with 70% ethanol (120 ul), air dried for 15 minutes in a fume hood, and resuspended in Qiagen EB (20 ul). DNA libraries were prepared using the Illumina Nextera DNA library Prep kit as per Baym M, et al (https://doi.org/10.1371/journal.pone.0131262) with modifications as follows. DNA was tagmented at 55C for 5 minutes. The adapter PCR program is: 72C for 3 minutes; 98C for 2 minutes 45 seconds; 8 cycles of 98C for 15 seconds, 62C for 30 seconds, 72C for 90 seconds; hold at 4C. An additional 4 cycles of reconditioning PCR were also performed. Libraries were pooled and size selected using AmpureXP to remove fragment <200 bp and minimize fragments >800 bp. Sequencing was performed on a Hiseq 4000 with 100 bp PE reads.

### Sequence processing

Code involved in the pipeline is deposited in github (https://github.com/setophaga/hybridswarm.alb.nas). Briefly we trimmed the reads with trimmomatic, with the following specification: -phred33 TRAILING:3 SLIDINGWINDOW:4:10 MINLEN:30. For some libraries, flies from up to four different species group were combined prior to DNA extraction, and reads were separated bioinformatically. In particular, we aligned the trimmed reads to the concatenated genome including the closest outgroup *D. kepuluana* genomic reference (26), as well as *D. pseudobscura* (GenBank: PRJNA596268), *D. virilis* (GenBank: PRJNA475270), and *D. athabasca* (27), with bwa (28). Individuals with less than 50,000 mapped reads were excluded from downstream analysis.

### Reference haplotypes

To construct parental ancestry haplotype reference, we used the existing high coverage sequencing data of alb03 line (2 males: DBMN30-16_S49_L008 and DBMN30-19_S51_L008, 1 female DBMN21-D_S4_L007) and nas00 line (DBCC035C4_S68_L008, DBMN21-B_S2_L007) (26). We aligned the reads to the same *D. kepluana* reference (see above) before genotyping with GATK 3.8. To determine *D. albomicans* vs. *D. nasuta* specific alleles, we calculated allele frequency within each species with VCFtools (29), and selected the sites with allele frequency difference greater than 0.3. Within *D. albomicans*-specific alleles, we determined whether a Muller CD alleles is neo-Y or neo-X specific following our previous study (19). Briefly, we regarded neo-sex chromosome-specific sites as those within Muller CD that are homozygous in females and heterozygous in males.

### Ancestry calling

Ancestry HMM (30) was used to infer local genomic ancestry among hybrids in the hybrid swarm experiment. The following setting was employed: *-a 2 0.5 0.5 -p 0 −3 0.5 -p 1 −3 0.5 -r 0.000005*. In particular, we assumed equal parental ancestry contributions, and recombination rate being 5×10^−6^, and estimated the generations before present in which the ancestry pulse occurred. We first found the ancestry informative SNPs with allele frequency difference > 0.3 between *D. nasuta* strain and *D. albomicans* strain. Then we prepared the input for ancestry HMM with a custom script. We defined 0 = *D. nasuta*, 0.5 = heterozygotes, 1 = *D. albomicans.* Ancestry genotype was only called if the maximum posterior probability > 0.9, when the maximum posterior probability genotype was assigned to the locus.

### Neo-sex chromosome haplotyping

Within Muller CD, we needed to delineate neo-X, neo-Y, versus *D. nasuta* ancestry blocks. To do so, we ran a secondary three ancestry types analysis within Ancestry HMM for sites within Muller CD, *-a 3 0.3325 0.3325 0.335 -p 0 −3 0.3325 -p 1 −3 0.3325 -p 2 −3 0.335 -r 0.000005*. A Muller CD haplotype was called if the maximum posterior probability > 0.9. For each individual, we estimated the haplotype proportion (proportion of sites of each haplotype across all the haplotype-informative sites). With this information, we could track the haplotype and genotype frequency over time. Codes in the pipeline is deposited in github (https://github.com/setophaga/hybridswarm.alb.nas).

### Genomic barriers to introgression

We identified genomic barriers to introgression based on the reduced ancestry junctions as well as the lack of admixture inferred from heterozygosity-ancestry relationship. Genomic barriers to introgression are expected to harbor reduced ancestry turnovers and suppression of admixture. We estimated ancestry turnover rate by the number of ancestry turnovers over the total ancestry informative sites in each Muller element. To test if there is difference in ancestry turnover rates, we employed Pearson’s Chi-square test with input being a contingency table of counts of ancestry junctions over total ancestry informative sites in each Muller element.

In addition, we considered genomic local admixture as a function of ancestry score (with 0 and 1 being pure *D. nasuta* and *D. albomicans,* respectively) and interspecific heterozygosity. Heterozygosity is expected to decay as admixture progresses. To control for pseudo-replication due to linkage among ancestry informative sites, we first identified genetic clusters within which the ancestry blocks tend to co-segregate among all the hybrids (31). Specifically, with R function *kmeans*, we iteratively incremented k (from k = 1) until 60% of the total variance was explained by between-clusters variance. For each K-means cluster, we calculated the barrier effect (the *γ* index) as a function of heterozygosity (*h*), the fraction of heterozygous ancestry-informative sites within each individual, and admixture proportion (*p*), the fraction of *D. albomicans* alleles across ancestry-informative sites.

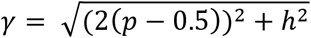

This index effectively represents the Euclidian distance between the position of each individual and the admixture maxima, where *p* = 0.5 and *h* = 0, in the triangle plot. The greater the barrier effect, the less admixture occurs within the genetic cluster, the greater distance the hybrids are from the point of admixture maxima. To compare the barrier effect *γ* among different Muller elements, we employed ANOVA followed by post-hoc paired-t-test with Bonferroni correction. Because the ancestral Y is largely pseudogenized and degraded, *γ* is not applicable for Muller A in males.

### Muller CD Introgression

To compare patterns of introgression in Muller CD versus the rest of the genome, we examined the ancestry proportion, inter-specific heterozygosity, linkage disequilibrium, and the genomic clines (32) of Muller CD relative to the rest of the genome.

For each K-means cluster (see above), we fit the Muller CD genomic cline (32) in which the mean ancestry of Muller CD clusters is the mean ancestry of all the K-means clusters genome-wide. In the course of admixture, local ancestry relative to genome-wide ancestry reflects the extent and direction of introgression of a genomic region. The genomic cline function *θ* = *p* + (2 (*p* - *p*^2^) × (*α* + (*β* (2*p*) - 1)), where *θ* is the local ancestry of Muller CD, and *p* is the genome-wide ancestry; *α* and *ß* represents the direction and extent of the barriers’ effect of the genomic region, respectively. If Muller CD introgressed similarly as the rest of the genome, *α* and *ß* should be zero. A significant barrier effect of Muller CD would be reflected by a positive *ß*. If there is disproportionally *D. albomicans* biased introgression within Muller CD, *α* should be significantly positive, whereas *D. nasuta* biased introgression corresponds to a negative *α* value. Under the neo-X advantage hypothesis, we expect a significantly positive *α* in hybrids.

Another representation of a barrier to introgression is linkage disequilibrium (LD). LD is expected to decay as admixture precedes, while genomic barriers to introgression would preserve high LD. If Muller CD serves as a genetic barrier to introgression, LD among K-means clusters within Muller CD would remain high relative to the rest of the genome. We tested whether the mean LD is different among Muller elements. Other genomic regions that harbor epistasis with genes in Muller CD would remain in high LD with Muller CD as well.

### Time series of haplotype and genotype frequencies

To understand the evolution and effect of the neo-sex chromosome, we tracked the dynamics of Muller CD ancestry (i.e. *D. nasuta* or *D. albomicans* neo-X or neo-Y) in a sex-specific manner over generations of hybridization. We modeled haplotype frequencies as time series and tested whether the time series of neo-X, neo-Y, or *D. nasuta* Muller CD haplotype frequency demonstrate autocorrelation against the null stationary model. We employed the rank von Neumann ratio test (33), with the *serialCorrelationTest* function in R. If no temporal autocorrelation was observed, we tested the deviation of the haplotype frequencies from the corresponding expected values based on 50:50 admixture. For females, the expected haplotype frequency of neo-X versus *D. nasuta* should be 0.5, while for males, the expected frequency of neo-X, neo-Y, and *D. nasuta* should be respectively, 0.25, 0.25, and 0.5, respectively. We have only included generations in which there were more than 5 females and 5 males.

In addition, we tested the pairing compatibility hypothesis in which the presence of the neo-X could facilitate the increase of neo-Y frequency due to problems in meiosis with the unfused *D. nasuta* Muller CD. If so, the neo-X frequency in females should be positively correlated with the neo-Y frequency in males, after controlling for temporal autocorrelation as the cofounding factor. We therefore used a partial Mantel test with mantel.partial function in R. Finally, we tested whether Muller CD segregation deviates from Hardy-Weinberg equilibrium by contrasting the expected genotype frequencies under random paring of haplotypes versus observed genotype frequencies with a Pearson’s Chi-squared test.

### Population genetic model of neo-sex chromosome evolution

To better understand the dynamical feedback between hybrid incompatibility, meiotic drive, and selective advantage of the neo-X and disadvantage of the neo-Y chromosome, we modeled the evolution of karyotype frequencies through time as a single-locus continuous time model with separate sexes. Parental individuals were assumed to mate at random. Here we denote *D. nasuta* X and Y chromosome and the unfused Muller CD as *N_x_* and *N_y_*; and the X and Y of *D. albomicans* as *A_x_* and *A_y_*. Due to genetic incompatibilities, hybrid zygotes (*N_x_A_x_, N_x_A_y_*, and *A_x_N_y_*) were subject to a reduction in absolute fitness of (1 − *ρ*). As a result of coevolution of meiotic drivers between the neo-X and suppressors on the neo-Y (but not the ancestral Y or unfused Muller CD), a meiotic driver on the neo-X chromosome results in killing of *D. nasuta* Y/Muller-CD sperm of heterozygous (neo-X, *D. nasuta* Y) with a probability of *μ_H_*. We also include two putative within-species meiotic drivers on the neo-X and *D. nasuta* X that respectively kill the neo-Y and the *D. nasuta* Y sperm with probability of *μ_A_* and *μ_N_*. Finally, we introduce a putative additive selected advantage to the neo-X of magnitude *s_x_* and an additive disadvantage of the neo-Y of size *s_y_*. The result is a system of seven coupled differential equations giving the frequency, *F*, of each karyotype. The derivation and expression of the differential equations can be found in the *Supplementary Mathematica* notebook. We analyze the karyotype frequency dynamics by first identifying the biologically valid equilibria of the dynamical system. We then analytically determine the local stability of these equilibria and finally use a numerical approach to determine the global stability of the equilibria given the initial conditions of the hybrid swarm.

### Fly crosses to estimate sex-ratio meiotic drive

5 virgin *D. albomicans* females were crossed to 7-10 *D. nasuta* males. F1 hybrid virgin females were backcrossed to either *D. albomicans* or *D. nasuta* males, and F1 hybrid males were backcrossed to either *D. albomicans* or *D. nasuta* virgin females. For each of these four backcrosses, three vials of 5 virgin females by 7-10 males were set up, and flies were were transferred to new vials every 2-3 days for 2 weeks. Once the adult offspring started emerging, offspring were sexed and counted everyday for up to ten days. Offspring counts from different vials and transfers of the same cross were then summed.

We estimate *D. albomicans* specific (*μ_A_*), *D. nasuta* specific (*μ_N_*), and heterospecific (*μ_H_*) meiotic drive with female and male counts with the following formula: (female counts – male counts)/female counts, as the fraction of Y gametes that were killed. We estimated *D. albomicans* and *D. nasuta* specific meiotic drive *μ_A_* and *μ_N_* from backcrossing F1 females (neo-X, *D. nasuta* Muller CD) respectively with *D. albomicans* males (neo-X, neo-Y) or *D. nasuta* males (*D. nasuta* Muller CD, *D. nasuta* Muller CD). To estimate *μ_H_*, we backcrossed F1 males (neo-X, *D. nasuta* Muller CD) with *D. albomicans* females (neo-X, neo-X) or *D. nasuta* females (*D. nasuta* Muller CD, *D. nasuta* Muller CD) and took the mean of the two backcrosses.

## Results

### Reduced introgression in Muller CD

Fig. 1B shows the inferred haplotypes of hybrids sampled over 62 generations of hybridization. There was extensive introgression and reshuffling of ancestral haplotypes in Muller A, B, and E, relative to Muller CD (**Fig. 1B**). Ancestry junction rates varied significantly among Muller elements (*χ*^2^ = 11.56, df = 3, *p* = 0.009), and the ancestry junction rate was reduced in Muller CD relative to other chromosomes (**Fig. 1–2**). The region of reduced ancestry turnover within Muller CD coincides with the two overlapping chromosomal inversions found in the strains used to generate the hybrid swarm (35) (**Fig. 2**).

**Fig. 2.**
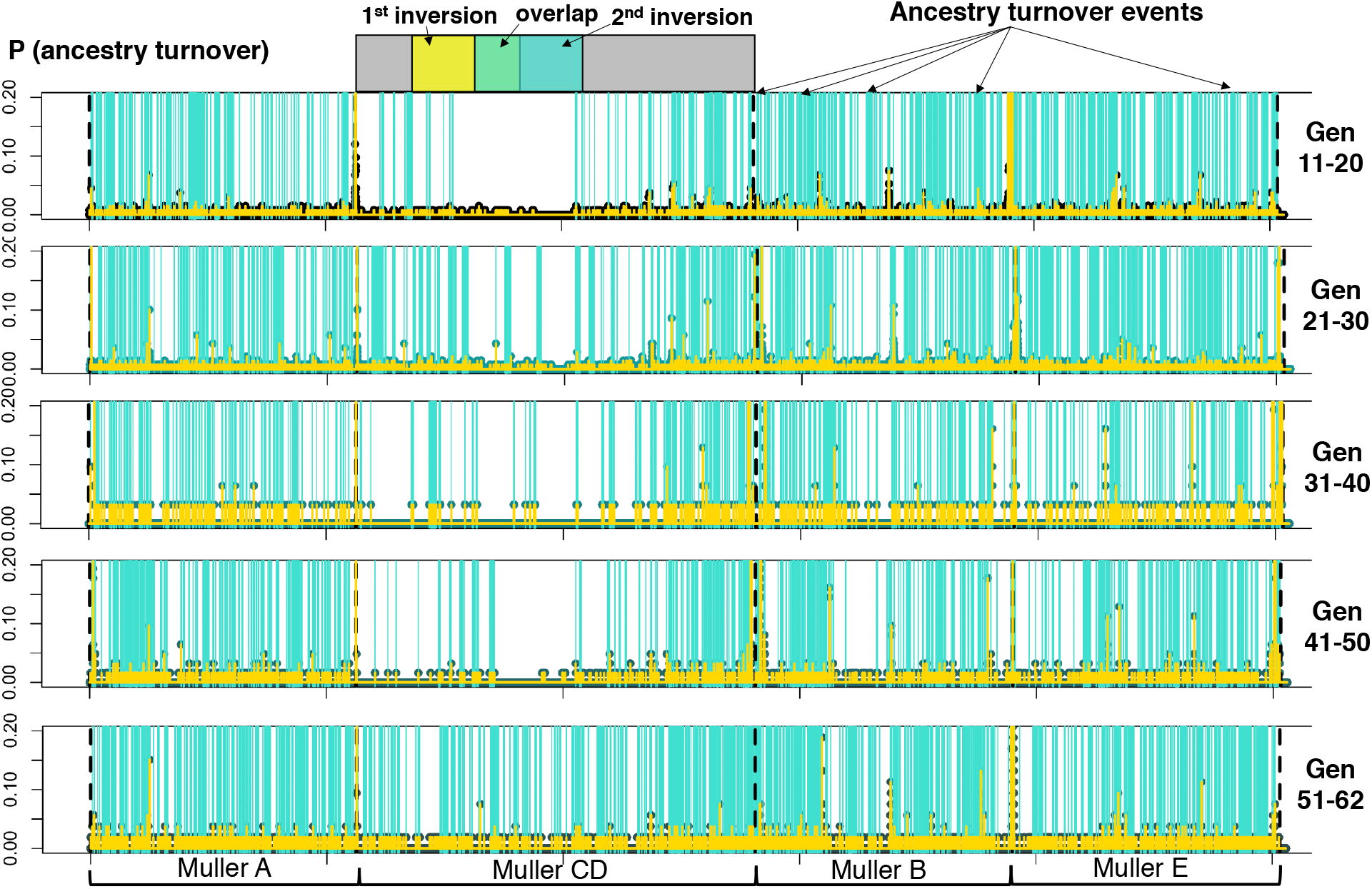
Incidences of ancestry turnovers during the course of hybrid swarm evolution. Each vertical turquoise line represents an ancestry junction. The yellow peaks represent the probabilities of ancestry turnovers across chromosomal positions. The reduced ancestry turnover rate in Muller CD corresponds to two pericentromeric overlapping inversions.

K-means clustering drastically reduces the number of ancestry-informative units in each Muller element and minimizes pseudo-replication (**Table S2**). Admixture proportions, which measure the fraction of *D. albomicans* alleles across each k-means cluster along each Muller element, were traced across generations in female versus male hybrids separately (**Fig. 3**). There was significant *D. albomicans*-biased introgression genome-wide in both females (mean= 0.65, t =15.70, *p* < 10^−15^) and males (mean = 0.59, t = 7.91, *p* < 10^−12^), while there was heterogeneity across Muller elements (**Fig. 3**).

**Fig. 3.**
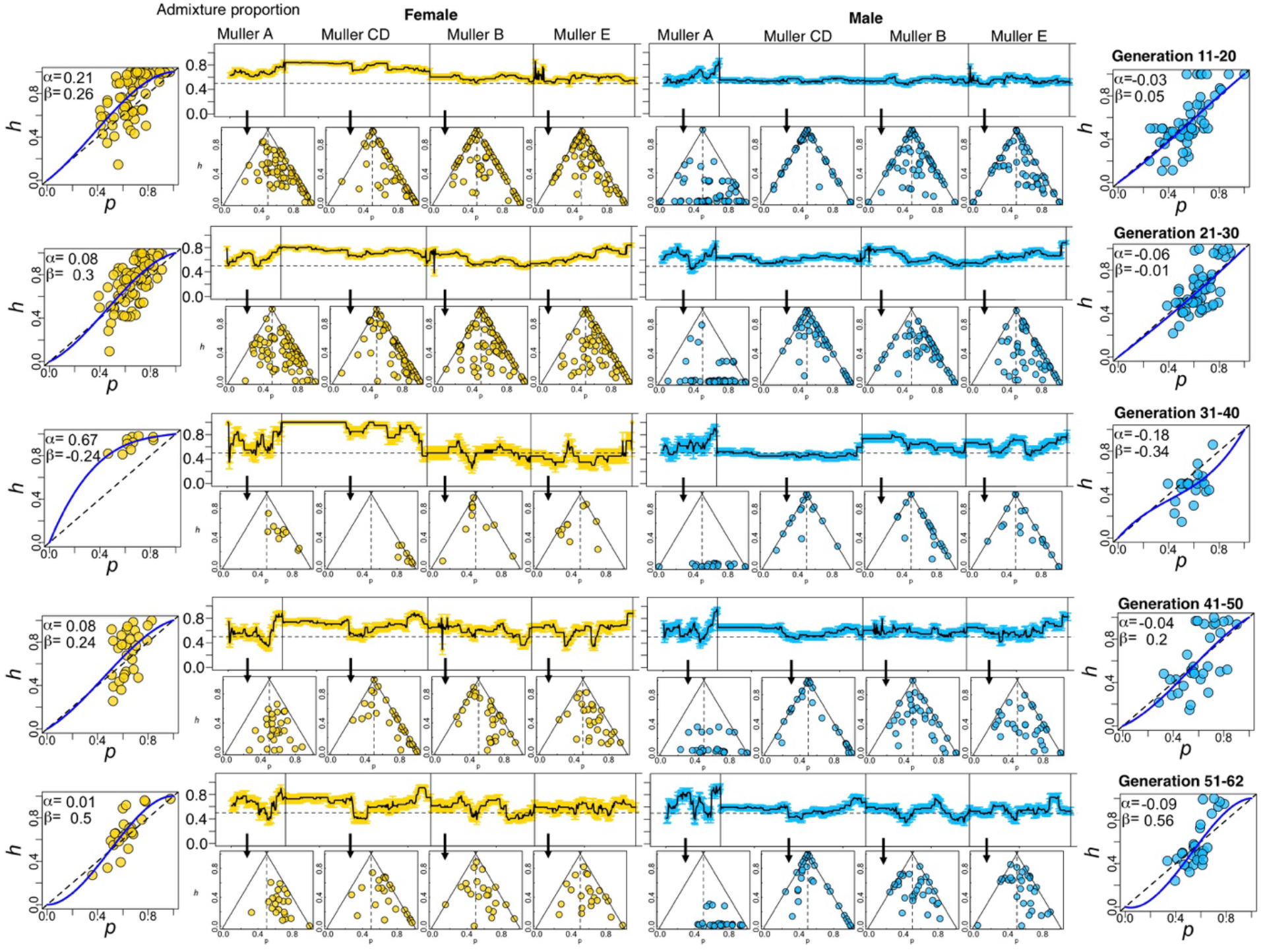
Patterns of introgression along chromosomes in female versus male hybrids sampled over generations. Each row represents introgression patterns in hybrids sampled from each generation interval: 11-20, 21-30, 31-40, 41-50, and 51-62, with females colored in yellow and males colored in blue. The ancestry proportion scan represents the mean (± SE) admixture proportion (pure *D. nasuta* = 0, pure *D. albomicans* = 1) of each K-means cluster in Muller A, CD, B, or E in females or males sampled across generations. The triangle plots present the relationship of mean heterospecific heterozygosity and ancestry proportion of each Muller elements. There is *D. albomicans*-biased introgression (admixture proportion > 0.5, horizontal dotted line; dots disproportionally shifted to the right arm of the triangle, alpha > 0) in Muller CD of females, but not for males. The genome clines represent Muller CD ancestry (θ) relative to genome-wide ancestry (p) in individuals sampled from each sex at each generation. The cline parameters *α* and *ß* represent the direction and extent of introgression. For *α*, positive values correspond to *D. albomicans*-biased introgression of Muller CD, whereas *D. nasuta*-biased introgression for negative values. Large positive *ß* corresponds to strong introgression barrier effect of Muller CD.

The barrier effect *γ*, which measures the reduction in admixture relative to the admixture maxima, was significantly elevated in Muller CD compared to other Muller elements in both sexes across all the generation intervals examined (**Fig. S1**; **Table S3**). The K-means clusters with the strongest barrier effect are associated with the overlapping inversion within Muller CD. Another measure of a barrier to introgression is linkage disequilibrium (LD). LD remained high after 62 generations of hybridization within Muller CD, while LD decays over the course of admixture in most of the genome (**Fig. 4; Fig. S4**). The mean pairwise LD was significantly longer within Muller CD than other Muller elements (*p* < 0.05; **Fig. S4**). In addition, some long-ranged LD was observed between Muller B and Muller E (**Fig. 4**). Consistent with elevated levels of LD, disproportionately less ancestry turnover events (0.83%) are detected in Muller CD, compared to Muller A (15.04%), Muller B (25.20%), and Muller E (58.92%).

**Fig. 4.**
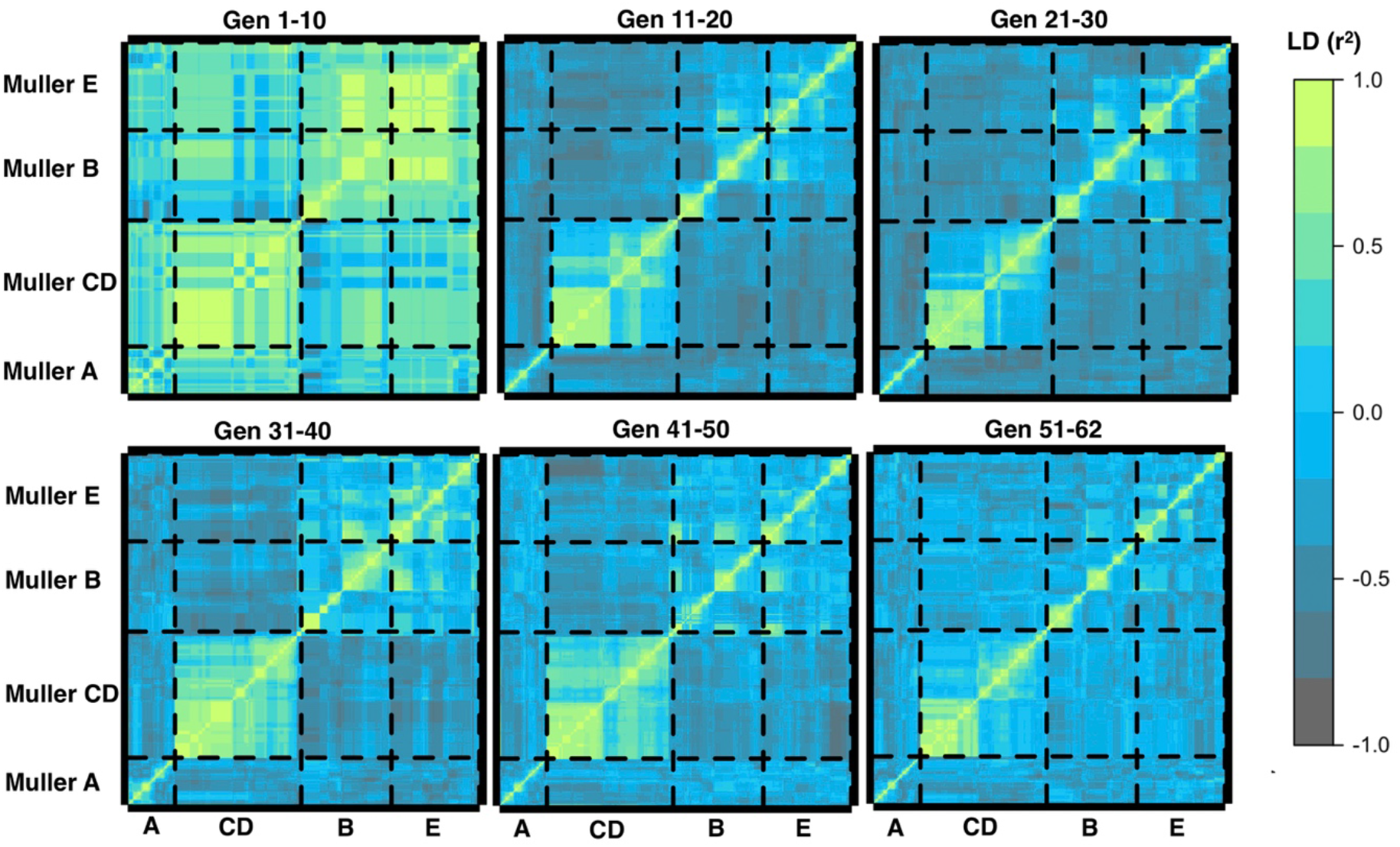
Linkage disequilibrium (*r^2^*) among K-means clusters across Muller elements. Genome-wide LD decays with admixture. However, LD remains high within Muller CD, and some regions in Muller B and E.

### Sex-dependent asymmetrical Introgression

Intriguingly, we find evidence for *D. albomicans*-biased introgression of Muller CD in females (admixture proportion > 0.5; **Fig. 3** yellow), but not in males. The genomic cline analysis revealed significantly positive *α* (*D. albomicans* biased introgression of Muller CD relative to genomic background) in generation 1-20 as well as 31-40 (**Table1**; **Fig. 3** yellow). In contrast, there was significantly negative *α* (*D. nasuta* biased introgression) in male hybrids sampled from generation 31-40 (**Table 1**; **Fig. 3** blue). Thus, the neo-sex chromosomes / Muller CD show sex-dependent, asymmetrical introgression during the course of admixture in our experimental hybrid swarms.

**Table 1.** Genomic cline parameter estimates of females and males in various generations.

| Generations | parameter | female estimate | female *p*-value | male estimate | male *p*-value |
| --- | --- | --- | --- | --- | --- |
| 11-20 | alpha | **0.21** | **0.01** | -0.03 | 0.55 |
|  | beta | 0.26 | 0.23 | 0.05 | 0.82 |
| 21-30 | alpha | 0.08 | 0.24 | -0.06 | 0.42 |
|  | beta | 0.30 | 0.12 | -0.01 | 0.97 |
| 31-40 | alpha | **0.67** | **< 10<sup>-5</sup>** | **-0.18** | **0.03** |
|  | beta | -0.24 | 0.22 | -0.34 | 0.31 |
| 41-50 | alpha | 0.08 | 0.60 | -0.04 | 0.66 |
|  | beta | 0.24 | 0.65 | 0.20 | 0.55 |
| 51-62 | alpha | 0.01 | 0.93 | -0.09 | 0.20 |
|  | beta | 0.50 | 0.13 | 0.56 | 0.08 |

### Muller CD haplotype frequency time series

No significant temporal autocorrelation was observed in the Muller CD haplotype frequency for neo-X, neo-Y, or *D. nasuta*, in females or male hybrids (*p* > 0.05), indicating the lack of directional change in haplotype frequency over the course of admixture. Different from the expectation of pairing compatibility, the residual of a linear model in which male neo-Y frequency is predicted by female neo-X frequency does not demonstrate temporal autocorrelation (*ρ* = −0.47, *p* = 0.22; **Fig. S3A**). We further tested whether the mean haplotype frequencies significantly deviated from expectation. There was significantly higher neo-X frequency (mean ± SD = 0.74 ± 0.11) in female hybrids than 0.5 (*p* < 10^−6^), which corresponds the significantly lower *D. nasuta* variant than expected. In contrast, the neo-Y haplotype frequency (mean ± SD = 0.12 ± 0.12) in male hybrids was significantly lower than expected value of 0.25 (t = − 4.38, *p* = 0.0005; **Fig. S3B**). There was higher than expected frequency of neo-X in male hybrids (t = 11.17, *p* < 10^−7^), while the *D. nasuta* variant did not significantly deviate from the 0.5 expectation (t = −1.15, *p* = 0.27).

Neo-X advantage is evident over generations of admixture in the hybrid population (**Fig. 5 AC**). Across generations, female hybrids predominantly carry (neo-X, neo-X) genotype (**Fig. 5A**), at a mean frequency of 0.55, which was significantly higher than neutral expected frequency of 0.25 (t = 6.28, *p* < 10^−4^). The mean frequency of (*D. nasuta* Muller CD, *D. nasuta* Muller CD) was 0.05, rarer than the expected mean frequency of 0.25 (t = −17.83, *p* < 10^−11^). The mean frequency of heterozygous (neo-X, *D. nasuta* Muller CD) was 0.36, which is significantly lower than neutral expectation of 0.5 (t = −3.4915, *p* = 0.003).

**Fig. 5.**
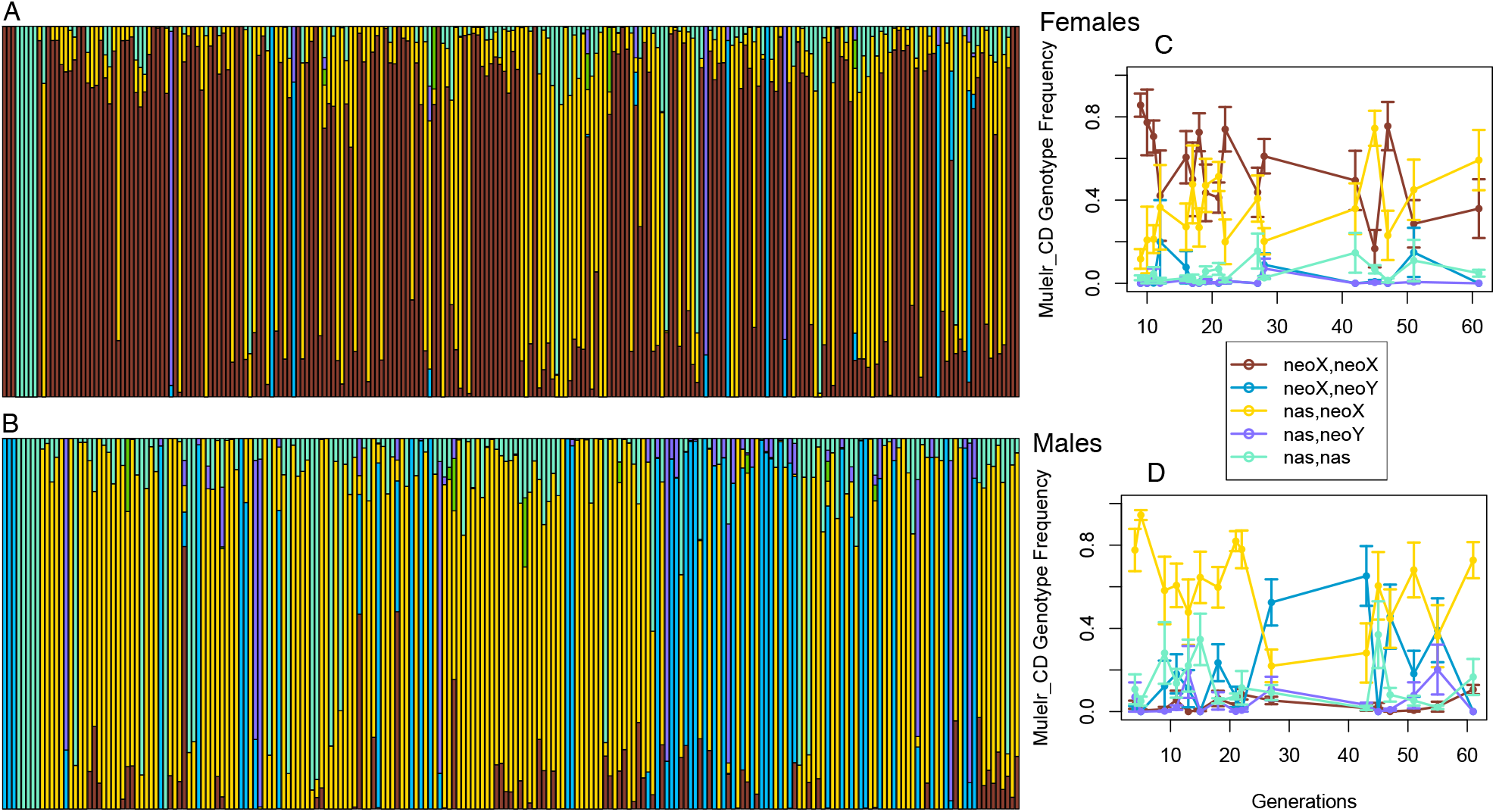
Sex-stage-dependent introgression asymmetry within Muller CD. Muller CD haplotype proportions in female (**A**) and male (**B**) hybrids sampled and ordered from generation 0 to 62. **C**, (neo-X, neo-X) was the predominant genotype in female throughout the generations sampled, but the heterozygous genotype (neo-X, *D. nasuta* Muller CD) and (neo-X, neo-Y) alternates in being the predominant male genotype (**D**). The error bars represent standard error of the mean.

In contrast, we observed male-specific heterosis (**Fig. 5 BD**), where the heterozygous genotype is more prevalent than homozygous genotypes. There was excessive heterozygous genotype of (neo-X, *D. nasuta* Muller CD) (mean frequency = 0.60) (t = 6.97, *p* < 10^−5^), but deficient (*D. nasuta* X, *D. nasuta* Muller CD) (mean frequency = 0.14) (t = − 4.06, *p* = 0.001) and (*D. nasuta* Muller CD, neo-Y) (mean frequency = 0.05) genotypes (t = −12.06, *p* < 10^−8^) than their expected 0.25 neutral frequency in male hybrids (**Fig. 5 D**). The (neo-X, neo-Y) frequency (mean = 0.18) did not significantly deviate from the expected value of 0.25 (*p* > 0.05) (**Fig. 5 D**).

### Population genetic model

The system of differential equation for the karyotype frequencies exhibits three equilibria, which are stable in at least a portion of the biological parameter space. Denoting equilibrium values with a hat:

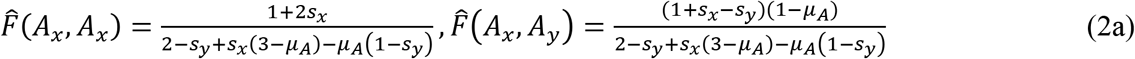

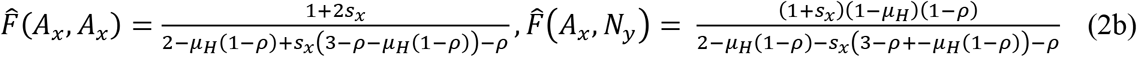

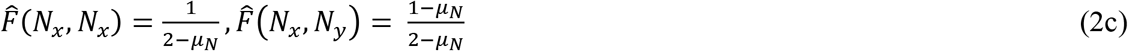

As the general conditions for the stability of these three equilibria are complex, we consider equilibrium stability and model dynamics for four specific sub-parameter cases *Case I, II, III, and IV* described below. To explore parameter space that could result in observed karyotypes in the end of the hybrid swarm experiment, we initialized the dynamics with even ancestry and sex ratio, which is consistent with the initialization of the hybrid swarm experiment.

We focus on Equilibrium 2b where (*A_x_*, *A_x_*) and (*A_x_*, *N_y_*) are the predominant karyotypes, because this equilibrium is the most consistent with the observed karyotype frequency at generation 62 of the hybrid swarm experiment. The temporal dynamics of the theoretical model exhibit no cyclic behaviour (all eigenvalues are real). Hence, while the karyotypes at generation 62 of the experiment are not yet fixed at this equilibrium the dynamical trend is constant with an approach to this fixed state. In all four parameter cases, we include selection favouring the neo-X (*s_x_* > 0) and incompatibility between heterospecific chromosomes (*ρ* > 0) as empirically established. The remaining parameters conditions for each of the four cases are described below.

In *Case I* we consider the effect of meiotic drive within *D. nasuta* (*μ_N_* ≥ 0) and/or heterospecific meiotic drive (*μ_H_* ≥ 0) in the absence of any form of selective disadvantage to *A_y_* (*s_y_* = 0, *μ_A_* = 0). The analytical local stability analysis reveals that under no such condition is the focal equilibrium (2b) stable. Hence, the observation of the empirical karyotype frequencies requires selection against *A_y_* carrying zygotes *s_y_* ≥ 0 and/or gametic selection against *A_y_* Via *D. albomicans*-specific meiotic drive *μ_A_* ≥ 0.

To examine when *D. albomicans* meiotic drive alone can facilitate the fixation of the *A_x_A_x_*/*A_x_N_y_* karyotype, in *Case II* we allow *μ_A_* > 0 setting the remaining parameters to 0 (*s_y_* = *μ_H_* = *μ_N_* = 0). The global stability of the equilibria given the initial conditions of the hybrid swarm experiment are shown in **Fig. 6B**, where the boundary between Equilibrium 2c (dotted) and 2b (solid yellow) is given by 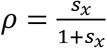 and the boundary between Equilibria 2a (fixation of *A_x_A_x_*/*A_x_A_y_*), 2b (fixation of *A_x_A_x_*/*A_x_N_y_*), and 2c (fixation of *N_x_N_x_*/*N_x_N_y_*) is determined numerically. The resulting parameter range over where the focal equilibrium (yellow region) is stable increases with an increasing strength of the *D. albomicans* meiotic drive *μ_A_*.

**Fig. 6.**
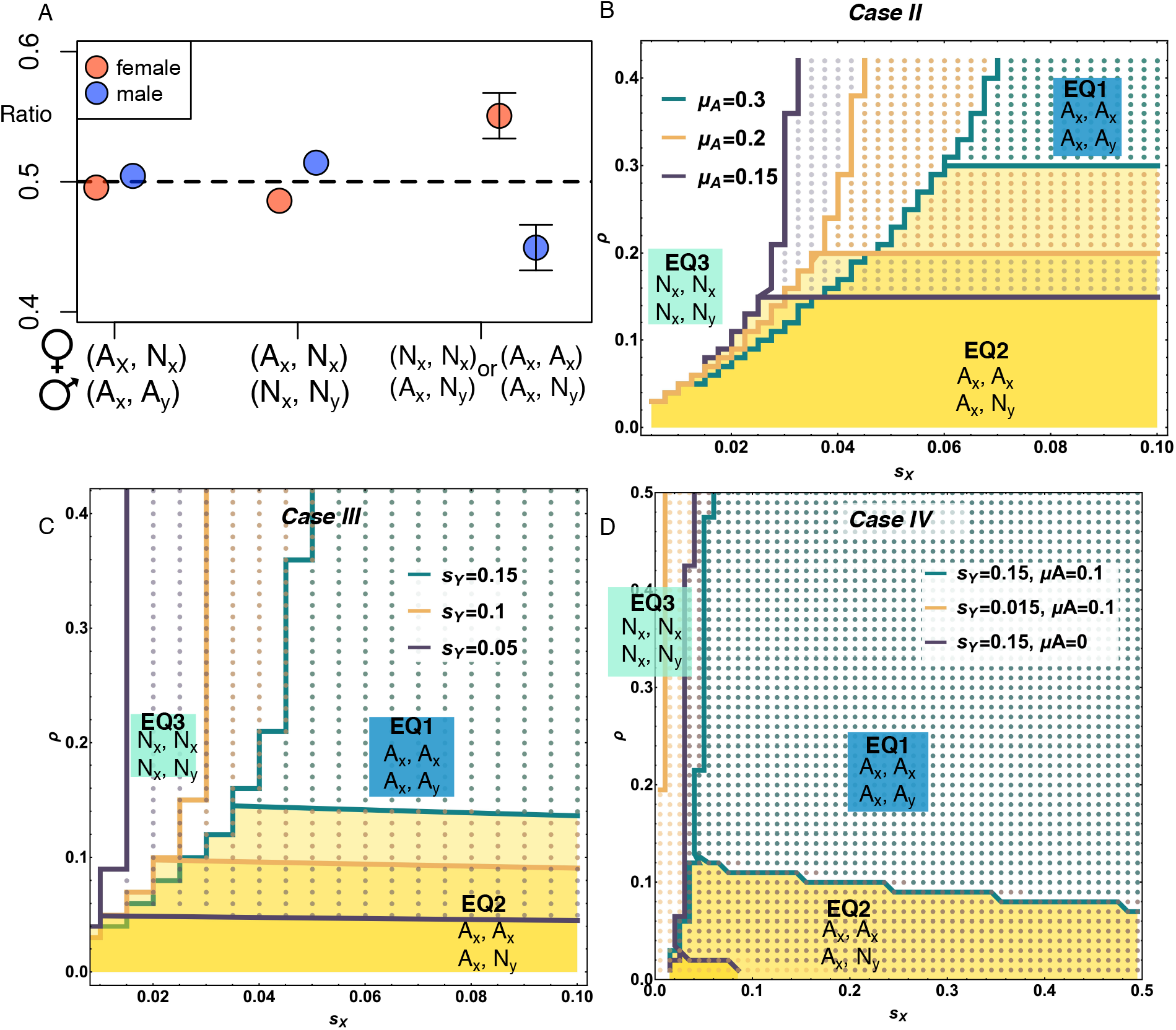
Parameter space in which the system reaches each of the three stable equilibria based on the initial condition of the hybrid swarm experiment. The parameter space in which equilibrium EQ1 (equation 2a-dotted region), EQ2 (equation 2b, yellow) and EQ3 (equation 2c-white) is reached given the initial conditions of the hybrid swarm experiment. **A**, Sex ratio assay reveals putative meiotic drive rate in conspecific (*μ_A_*, *μ_N_*) and heterospecific (*μ_H_*) conditions. Shown are inferred sex ratios in backcrosses, where female genotypes were listed above male genotypes. Conspecific meiotic drive rate *μ_A_* and *μ_N_* are respectively inferred from an (A_x_, N_x_) x (A_x_, A_y_) and (A_x_, N_x_) x (N_x_, N_y_) cross. No significantly sex ratio distortion is observed within *D. albomicans* or *D. nasuta*. However, there was significant sex ratio distortion between species (heterospecific meiotic drive *μ_H_*), which is estimated to be 0.12. **B**, In the population genetic model we fixed meiotic drive rates based on sex ratio estimates, where *μ_A_* = 0, *μ_H_* = 0, *μ_N_* = 0, *s_y_* > 0. The conditions for the hybrid swarm to reach EQ2, with (A_x_, A_y_) and (A_x_, N_y_) genotypes, is delineated in the yellow shades in the space of *ρ*, *s_X_*, and *μ_A_*. **C**, For *Case III*, where *ρ* > 0, *s_X_* > 0, *s_Y_* > 0 and *μ_A_* = *μ_N_* = *μ_H_* = 0, to reach EQ2, as observed in the hybrid swarm, requires strong *s_Y_* and low *ρ*. **D,** Result for *Case IV*, where *ρ* > 0, *s_X_* > 0, *s_Y_* > 0 and *μ_A_* > 0, *μ_N_* = 0, *μ_H_* = 0.12. To reach EQ2 in this case requires selection against the neo-Y, moderate selection for the neo-X, and weak hybrid incompatibility.

In contrast, in *Case III* we consider if and when the focal equilibrium is stable given only selection against *A*_*y*_ carrying zyogotes (*s_y_* > 0, *μ_A_* = *μ_H_* = *μ_N_* = 0). The resulting global stability analysis is shown in **Fig. 6B**. As with *Case II*, the boundary between Equilibrium 2a and 2b/2c is determined numerically and the boundary between 2b (yellow) and 2c (dotted) is given by 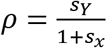. The parameter range under which the focal equilibrium is stable is relatively small, requiring *s_y_* to be large.

Direct application of *Case II* and *III* to the empirical system is limited as both assume *μ_H_* = 0 whereas the empirical estimates of *μ_H_* = 0.12 (**Fig. 6A**). While we are not able to examine the case of *μ_H_* ≠ 0 in general, in *Case IV* we examine three specific parameter combinations of *μ_A_* and *s_y_*, given *μ_H_* = 0.12. Overall, we find that *μ_H_* hinders the fixation of the *A_x_A_x_*/*A_x_N_y_* karyotypes (**Fig. 6D**). For this focal equilibrium to be stable across a substantial region of parameter space requires the presence of both strong zygotic selection *s_y_* and gametic selection *μ_A_* against *A_y_*.

## Discussion

By tracking *D. albomicans* - *D. nasuta* ancestry in hybrid swarms over many generations, we revealed the conflicting roles of neo-sex chromosome evolution in introgression: overall, the neo-sex chromosomes serve as sex-dependent asymmetrical introgression barriers between species. Limited introgression was observed within the overlapping paracentric inversions located on the neo-sex chromosome / Muller-CD. Female-specific *D. albomicans* biased introgression of the neo-X chromosome is consistent with the relative fitness advantage of neo-X versus the unfused *D. nasuta* type (Fig. 5C) and neo-X meiotic drive. Despite the fact that the neo-Y is the more compatible pairing partner of neo-X relative to the *D. nasuta* unfused type, we did not observe neo-X-facilitated neo-Y introgression. Instead, we observed male-specific heterosis (Fig. 5D). Theory suggests such conflicting sex-dependent pattern of introgression relies on either *D. albomicans* specific meiotic drive and neo-Y disadvantage in concert with neo-X advantage and heterospecific chromosome pairing disadvantage. Sex ratio assays revealed interspecific sex chromosome drive (**Fig. 6A**), and previous sequence and expression analysis showed that the neo-Y chromosome shows moderate levels of degeneration, and may thus be selected against in hybrid males (19). Our population model suggests that the interplay among meiotic drive, neo-Y degeneration disadvantage, neo-X advantage, and pairing incompatibility can account for the sex-dependent asymmetrical introgression barrier effect of the neo-sex chromosomes.

### Introgression barrier

Sex chromosomes are increasingly recognized as barriers of introgression across a diverse group of organisms (6–8, 36, 37). However, the mechanism of such a barrier effect is not well-understood. By tracking the behavior of newly-formed sex chromosomes at an incomplete species boundary, we dissected barrier effects of sex chromosome evolution on introgression. We observed a strong introgression barrier effect within the recently formed neo-sex chromosome (**Fig. 1**), which is partly explained by two overlapping inversions on Muller CD (**Fig. 2**). In addition, pairing incompatibility between the *D. albomicans* fused and *D. nasuta* unfused genotypes (21) may further explain the reduced introgression in Muller CD. After the fusion of the sex chromosome and autosome forming the neo-sex chromosome, the neo-sex chromosomes may accumulate sexual antagonistic loci (38, 39), which can also prevent introgression of heterospecific variants.

### Sex-dependent asymmetrical introgression

In addition to the barrier effect, we also observed sex-dependent asymmetrical introgression associated with the neo-sex chromosome (**Fig. 3**). In females, there was *D. albomicans* biased introgression, whereas there was heterosis in males (**Fig. 5D**). The fused *D. albomicans* neo-X chromosome is thought to be advantageous over the unfused primitive haplotype (21). For the neo-X chromosome to increase to high frequency, it has to overcome meiotic structural incompatibility with its unfused homolog. Female *D. albomicans* biased neo-X introgression (**Fig. 5**) supports neo-X advantage, since advantageous parental haplotypes tend to dominate hybrid genomes (40). The neo-X chromosome may be preferentially transmitted to the next generation over the unfused Muller CD, by hijacking the asymmetric divisions of female meiosis (female meiotic drive)(41). We used sequencing of a large pool of backcross progeny embryos to test for deviations from Mendelian segregation (following ref. 42), but found no evidence of meiotic drive in female F1 hybrids (**Fig. S5**). This suggests that the selective advantage of the neo-X is not due to some conflict during female meiosis.

However, neo-X advantages and/or pairing incompatibility does not explain heterosis in males (**Fig. 5D**). Surprisingly, the heterozygous (neo-X, *D. nasuta* Muller CD) genotype occurred more frequently than the neo-X and neo-Y combination in the beginning and the end of the hybrid swarm experiment (**Fig. S3**). Notably, the heterosis in the heterogametic sex is opposite to Haldane’s Rule.

Population genetic modeling suggests that the interplay among neo-Y disadvantage and meiotic drive in *D. albomicans*, combined with neo-X advantage and heterospecific pairing incompatibility can explain the observed male-specific heterosis and female-specific *D. albomicans* introgression in Muller CD (**Fig. 5–6**). With moderate meiotic drive and selective disadvantage of neo-Y, at moderate pairing incompatibility and selective advantage of neo-X, the system could reach the (neo-X, *D. nasuta* Muller CD) equilibrium in males. Meiotic drive alleles have been characterized in *D. albomicans* Muller CD with QTL mapping (24), in addition to neo-X advantage (21), and chromosomal pairing incompatibility between *D. nasuta* Muller CD and the fused type (21, 22). However, in the parental strains involved in the hybrid swarm experiment, we did not observe intraspecific meiotic drive in *D. albomicans* or *D. nasuta*, but there was low to moderate level of heterospecific meiotic drive (~12%, female bias). Patterns of molecular evolution and gene expression on the neo-Y of 15112-1751.03, the *D. albomicans* parental strain we used, revealed moderate levels of degeneration, with dozens of neo-Y-linked genes showing stop codons and frameshift mutations and reduced gene expression (19). This supports selection against the neo-Y in hybrid males (s_Y_). Our population model (**Fig. 6D**) suggests that under the initial condition of the experiment and estimations of meiotic drive rates, selection against the neo-Y (s_Y_ >0) is required to observe excessive (neo-X, *D. nasuta Muller CD*) combination, the male-specific Muller CD heterosis.

### Alternative oscillation of (neo-X, D. nasuta Muller CD) and (neo-X, neo-Y)

The most abundant Muller CD genotypes in males were (neo-X, *D. nasuta* Muller CD). However, an exception occurred between generation 30 and 40, when (neo-X, neo-Y) became more abundant (**Fig. 5D**); such oscillation suggests that the system might not have reached equilibrium. The low frequency of (*D. nasuta* X, *D. nasuta* Muller CD) throughout the experiment suggests that neo-X advantage could be strong. When neo-X increases in frequency in females, in males (neo-X, *D. nasuta* Muller CD) became more prevalent, instead of (neo-X, neo-Y). The neo-Y haplotypic frequency in males is less associated with neo-X frequency in females than *D. nasuta* unfused haplotype, which is likely due to the opposite directions of selection in neo-X versus neo-Y. Towards the last generation, males were mostly (neo-X, *D. nasuta* Muller CD) while other genotypes were almost entirely lost. This is similar to the EQ2 condition, which the system is more likely to arrive when there is selective advantage of neo-X, selective disadvantage of neo-Y, *D. albomicans* meiotic drive, and pairing incompatibility (**Fig. 6**).

## Conclusion

Here we characterized and dissected sex-dependent asymmetrical barriers to introgression in the early stage of divergence that are affected by the evolution of neo-sex chromosomes. Such complex genomic barrier effect can be explained by the interplay of neo-X advantage and neo-Y degenerative disadvantage, chromosomal pairing incompatibility, and meiotic drive within the neo-sex chromosome.

## Supplementary Figures

**Figure S1.**
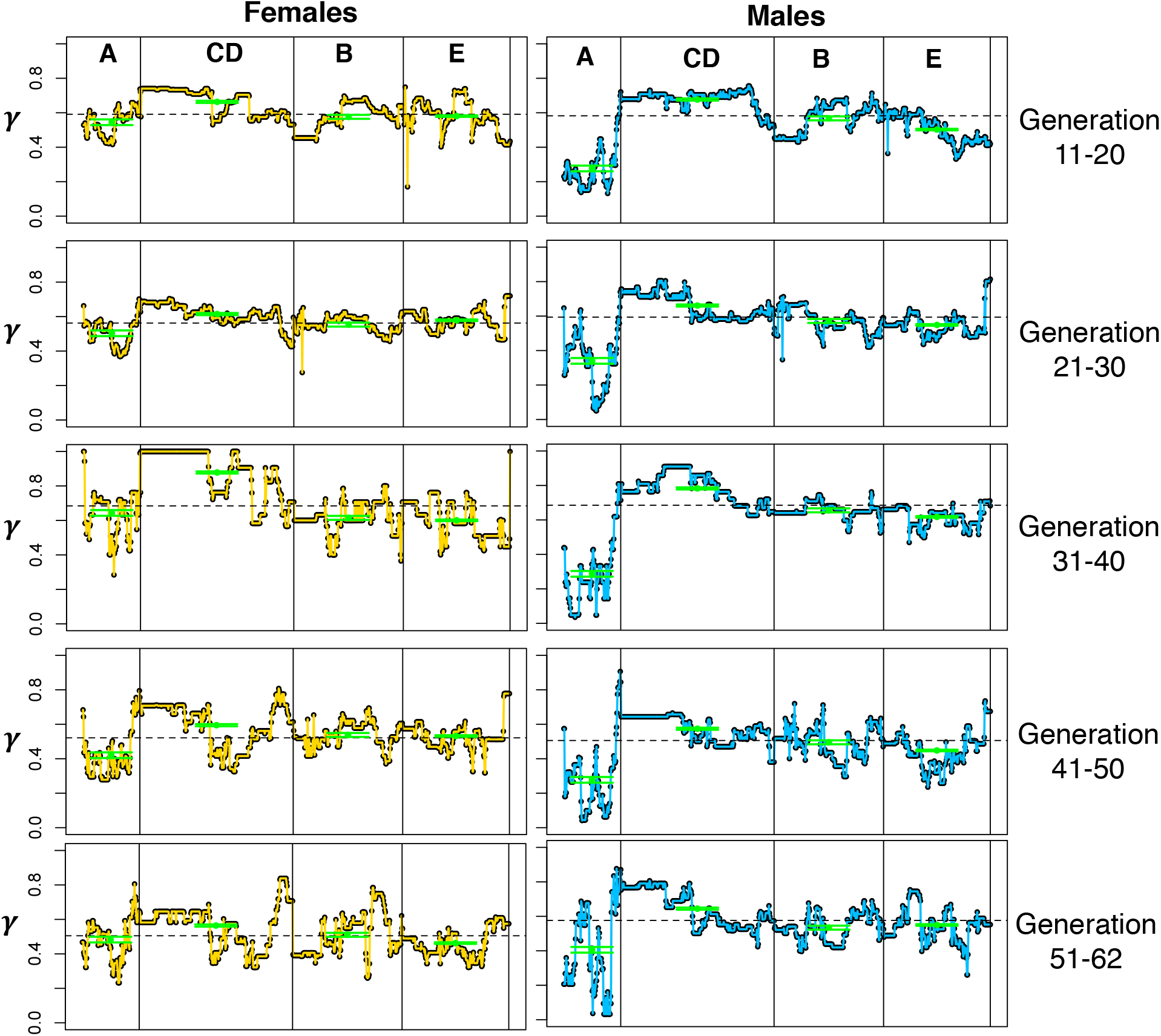
Barrier effect *γ* across chromosomes over generations. The green dot with arrow bars represent the mean (SE) of *γ* in each Muller element. There was significantly greater barrier effect in Muller CD than other Muller elements in both sexes across generations. The region of *γ* elevation is associated with overlapping inversions within Muller CD.

**Fig. S2.**
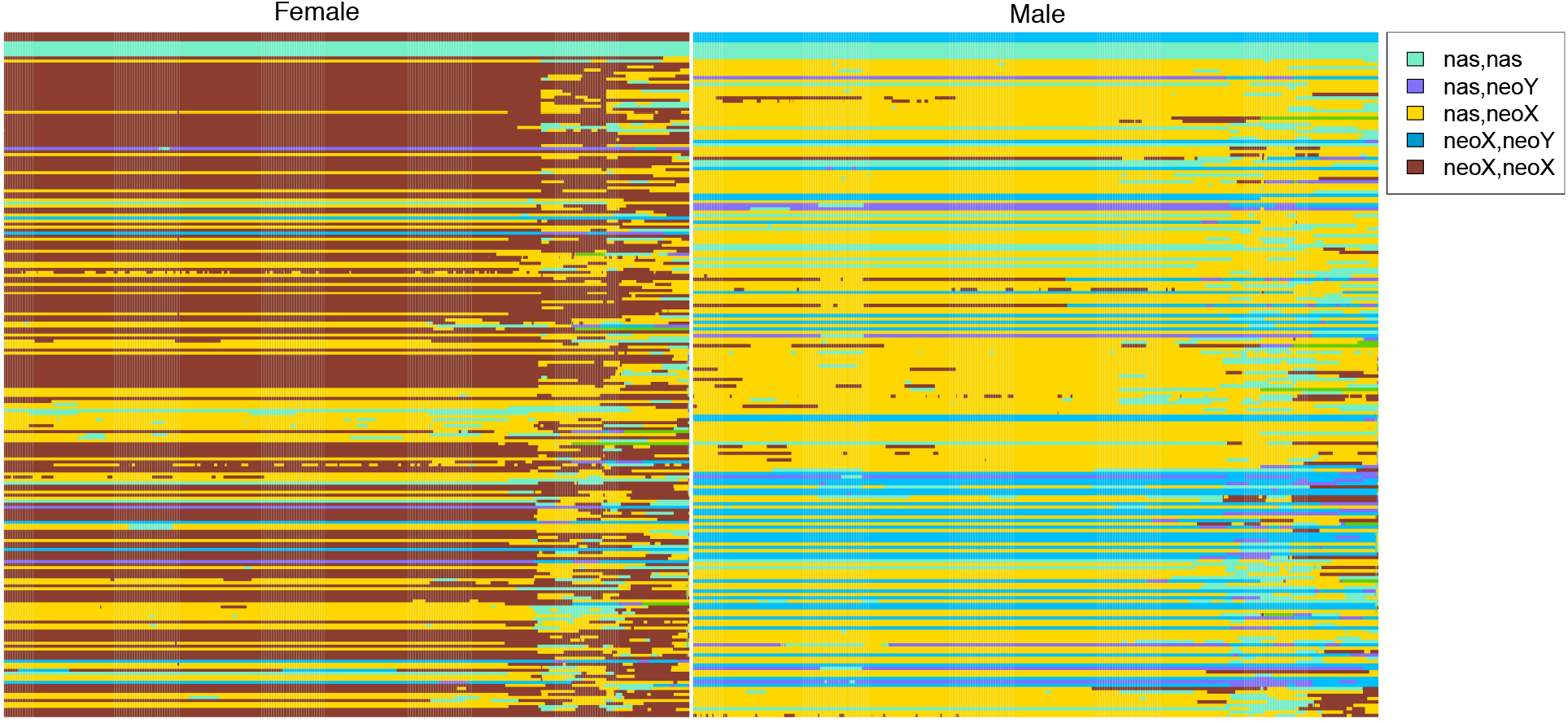
Karyotypes of Muller CD among hybrid females (left) and males (right). Each row represents the genotype of Muller CD of a hybrid, ordered from generation 0 to 62.

**Fig. S3.**
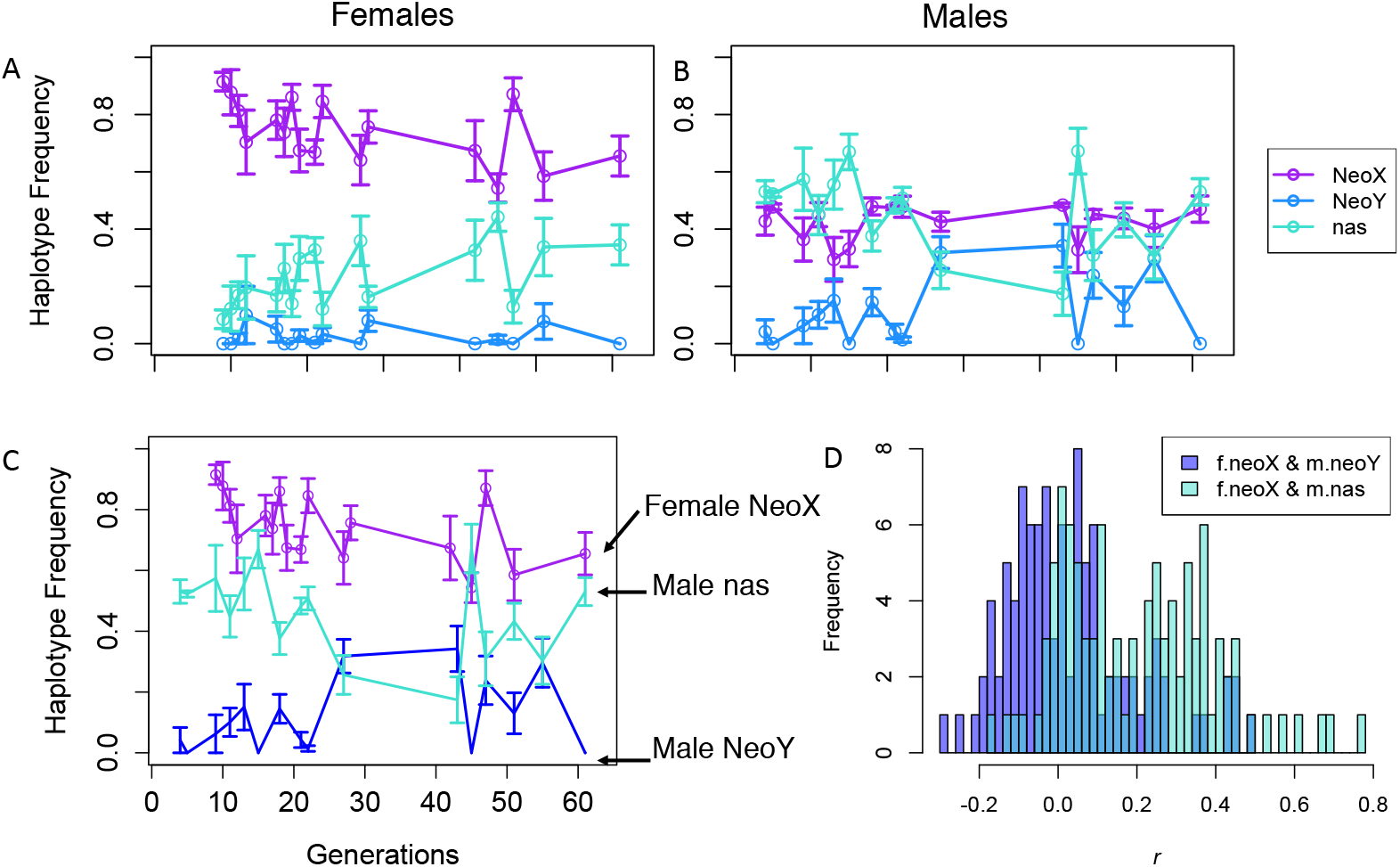
Time series of frequency of female neo-X, *D. nasuta* Muller CD, and male neo-Y. A, The frequency of neo-X is higher than 0.5 (*p* < 10^−6^) among female hybrids. B, Among male hybrids, the frequency of neo-Y oscillates below the expected 0.25 (*p* = 0.0005), and the frequency of *D. nasuta* Muller CD does not significantly deviate from expected value of 0.5 (*p* = 0.27). **C**, The oscillation of female neo-X is not positively correlated with male neo-Y as expected under pairing incompatibility prediction. **D**, Histograms of the partial mantel correlation coefficient (r) between female neo-X and male neo-Y, versus r between female neo-X and male *D. nasuta* Muller CD. Surprisingly, the latter was significantly shifted to the right (*p* < 0.05).

**Fig. S4.**
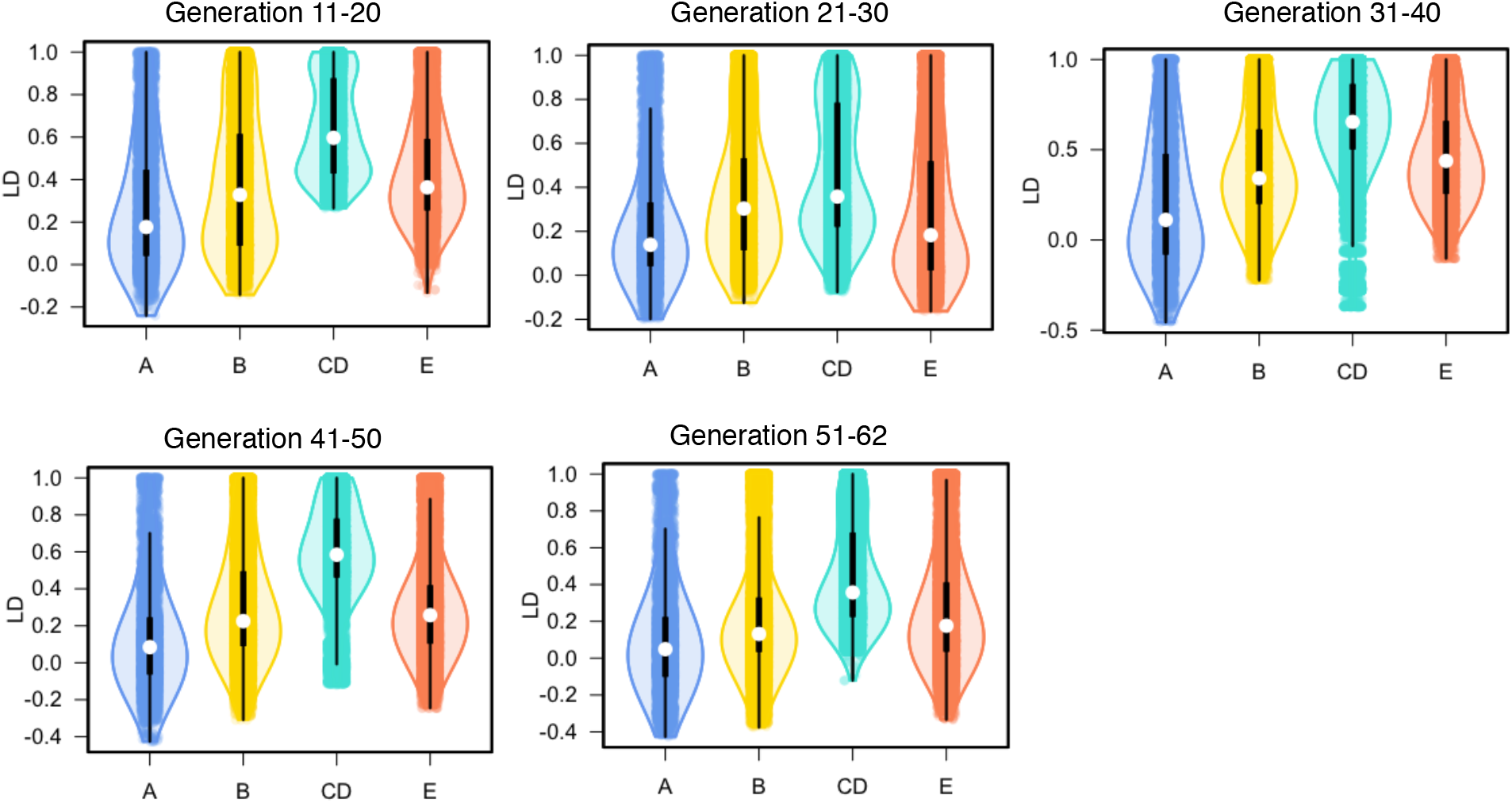
LD difference among Muller elements.

**Fig. S5.**
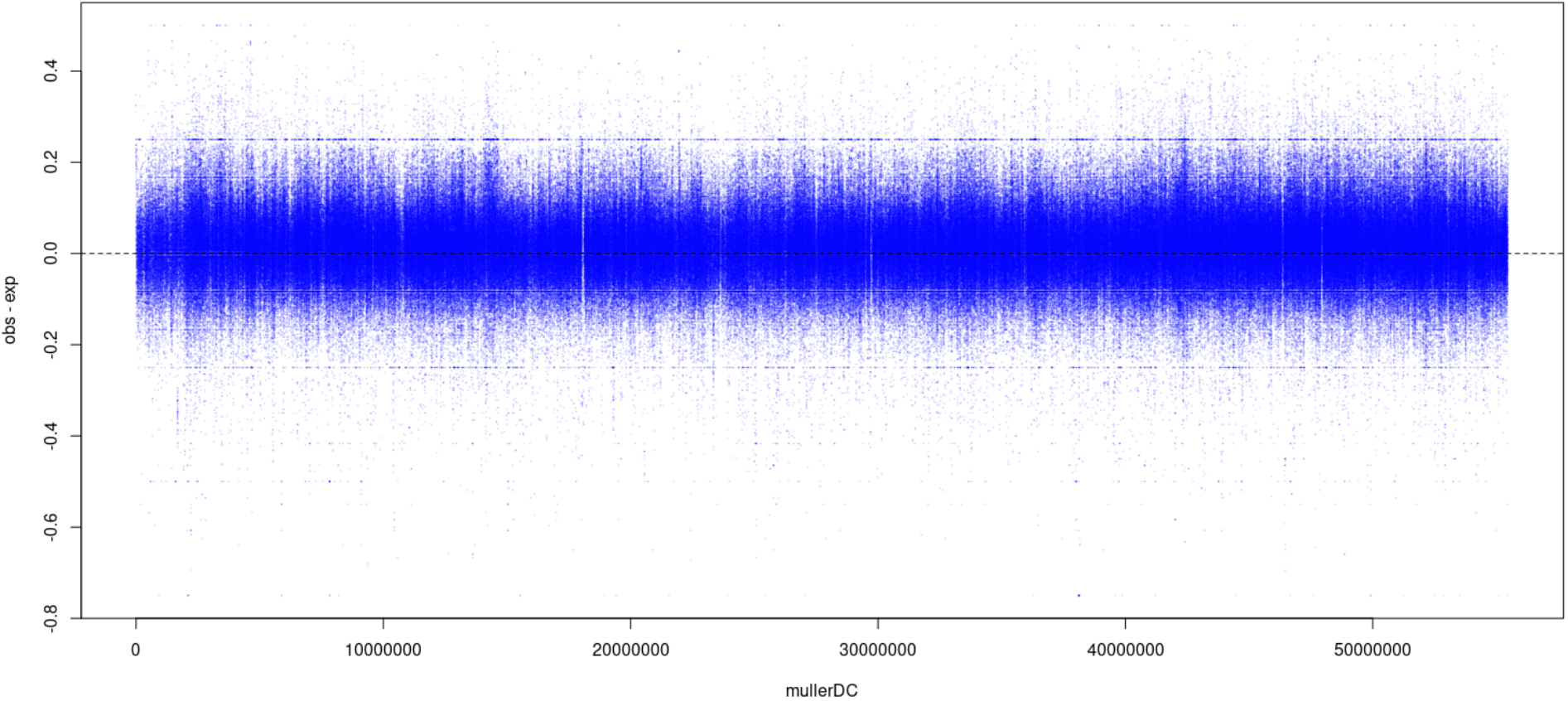
Mendelian segregation of Muller DC showing no evidence of meiotic drive in female hybrids. In the F1 hybrid females between *D. albomicans* and *D. nasuta,* the metacentric neo-X chromosome (Muller A fused with Muller CD) of *D. albomicans* form a tri-valent with the two acrocentric chromosomes (Muller A and unfused Muller DC) of *D. nasuta*. These F_1_ hybrids were mated to *D. albomicans* males and over 5000 3-4hr embryos were collected, pooled and sequenced. To infer deviation from expected Mendelian segregation, we first estimated the allele frequency of the embryo pool at polymorphic sites fixed between species. The observed allele frequency is subtracted from the expected Mendelian frequency at each site (blue point). While meiotic drive will cause broad and systematic deviation between the observed and expected frequencies, the pool show no such deviations.

**Table S1.** Number of flies sequenced in each generation of the hybrid swarm with sex information.

| Generation | 3 | 4 | 5 | 8 | 9 | 10 | 11 | 12 | 13 | 15 | 16 | 17 | 18 | 19 |
| --- | --- | --- | --- | --- | --- | --- | --- | --- | --- | --- | --- | --- | --- | --- |
| Female | 0 | 1 | 0 | 0 | 17 | 5 | 27 | 5 | 0 | 0 | 11 | 5 | 16 | 9 |
| Male | 1 | 12 | 5 | 2 | 8 | 1 | 16 | 0 | 9 | 11 | 2 | 0 | 20 | 4 |
| Generation | 21 | 22 | 27 | 28 | 42 | 43 | 45 | 47 | 48 | 51 | 55 | 61 | 62 |  |
| Female | 34 | 8 | 13 | 25 | 10 | 0 | 9 | 9 | 0 | 8 | 4 | 6 | 2 |  |
| Male | 28 | 10 | 18 | 0 | 3 | 10 | 6 | 10 | 1 | 11 | 10 | 9 | 0 |  |

**Table S2.** Numbers of ancestry informative sites and k-means clusters across each Muller element.

|  | Muller A | Muller B | Muller CD | Muller E | Muller F |
| --- | --- | --- | --- | --- | --- |
| Ancestry-informative sites | 112,545 | 108,773 | 168,676 | 113,222 | 2,648 |
| K-means clusters | 123 | 235 | 329 | 229 | NA |

**Table S3.** Difference in Barrier effects (*γ*) among chromosomes in female versus male hybrids sampled across generations. Within each generation interval for each sex, the ANOVA F value along with *p*-values were shown along with *p*-values for pairwise T tests with Bonferroni correction.

|  |  | Females |  |  |  |  | Males |  |  |  |  |
| --- | --- | --- | --- | --- | --- | --- | --- | --- | --- | --- | --- |
| Muller elements | | Muller A | Muller B | Muller CD | F | $p$-value | Muller A | Muller B | Muller CD | F | $p$-value |
| **Gen 10-21** | Muller A |  |  |  |  |  |  |  |  |  |  |
| | Muller B | 0.003 | | | 97.9 | $< 10^{-15}$ | $< 10^{-15}$ | | | 785.48 | $< 10^{-15}$ |
| | Muller CD | $< 10^{-15}$ | $< 10^{-15}$ | | | | $< 10^{-15}$ | $< 10^{-15}$ | | | |
| | Muller E | 0.0004 | 1 | $< 10^{-15}$ | | | $< 10^{-15}$ | $< 10^{-15}$ | $< 10^{-15}$ | | |
| **Gen 21-31** | Muller A |  |  |  |  |  |  |  |  |  |  |
| | Muller B | $< 10^{-12}$ | | | 118.7 | $< 10^{-15}$ | $< 10^{-15}$ | | | 387.49 | $< 10^{-15}$ |
| | Muller CD | $< 10^{-15}$ | $< 10^{-15}$ | | | | $< 10^{-15}$ | $< 10^{-15}$ | | | |
| | Muller E | $< 10^{-15}$ | $< 10^{-4}$ | $< 10^{-10}$ | | | $< 10^{-15}$ | 0.035 | $< 10^{-15}$ | | |
| **Gen 31-40** | Muller A |  |  |  |  |  |  |  |  |  |  |
| | Muller B | 0.19 | | | 343.56 | $< 10^{-15}$ | $< 10^{-12}$ | | | 847.74 | $< 10^{-15}$ |
| | Muller CD | $< 10^{-15}$ | $< 10^{-15}$ | | | | $< 10^{-15}$ | $< 10^{-15}$ | | | |
| | Muller E | 0.008 | 1 | $< 10^{-15}$ | | | $< 10^{-15}$ | $< 10^{-4}$ | $< 10^{-10}$ | | |
| **Gen 41-50** | Muller A |  |  |  |  |  |  |  |  |  |  |
| | Muller B | $< 10^{-15}$ | | | 83.13 | $< 10^{-15}$ | $< 10^{-15}$ | | | 233.14 | $< 10^{-15}$ |
| | Muller CD | $< 10^{-15}$ | $< 10^{-8}$ | | | | $< 10^{-15}$ | $< 10^{-15}$ | | | |
| | Muller E | $< 10^{-15}$ | 1 | $< 10^{-10}$ | | | $< 10^{-15}$ | $< 10^{-4}$ | $< 10^{-10}$ | | |
| **Gen 51-62** | Muller A |  |  |  |  |  |  |  |  |  |  |
| | Muller B | 0.16 | | | 97.9 | $< 10^{-15}$ | $< 10^{-15}$ | | | 115.33 | $< 10^{-15}$ |
| | Muller CD | $< 10^{-10}$ | $< 10^{-6}$ | | | | $< 10^{-15}$ | $< 10^{-15}$ | | | |
| | Muller E | 0.62 | $< 10^{-4}$ | $< 10^{-15}$ | | | $< 10^{-15}$ | 1 | $< 10^{-10}$ | | |

**Table S4.**
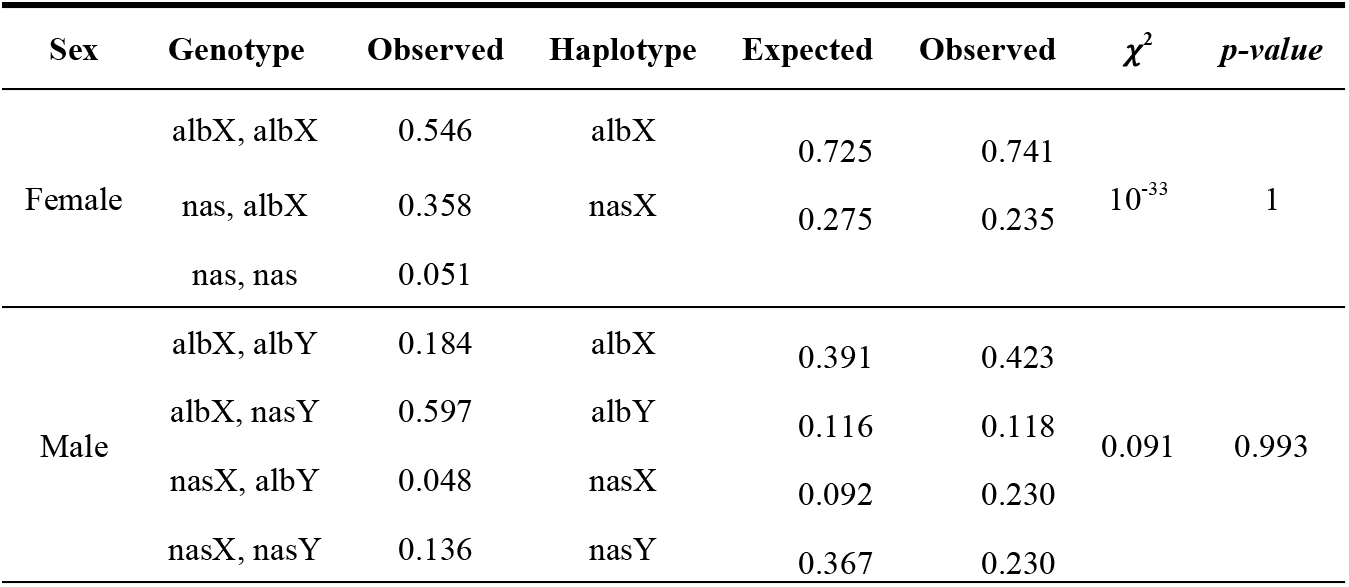
Muller CD haplotype frequency did not deviate from Hardy-Weinberg equilibrium

| Sex | Genotype | Observed | Haplotype | Expected | Observed | $\chi^2$ | *p-value* |
| --- | --- | --- | --- | --- | --- | --- | --- |
| Female | albX, albX | 0.546 | albX | 0.725 | 0.741 | $10^{-33}$ | 1 |
|  | nas, albX | 0.358 | nasX | 0.275 | 0.235 |  |  |
|  | nas, nas | 0.051 |  |  |  |  |  |
| Male | albX, albY | 0.184 | albX | 0.391 | 0.423 | 0.091 | 0.993 |
|  | albX, nasY | 0.597 | albY | 0.116 | 0.118 |  |  |
|  | nasX, albY | 0.048 | nasX | 0.092 | 0.230 |  |  |
|  | nasX, nasY | 0.136 | nasY | 0.367 | 0.230 |  |  |

